# Hybrid optical gating for long-term 3D time-lapse imaging of the beating embryonic zebrafish heart

**DOI:** 10.1101/526830

**Authors:** Jonathan M. Taylor, Carl J. Nelson, Finnius A. Bruton, Aryan K. Baghbadrani, Charlotte Buckley, Carl S. Tucker, John J. Mullins, Martin A. Denvir

## Abstract

Three-dimensional fluorescence time-lapse imaging of structural, cellular and sub-cellular processes in the beating heart is an increasingly achievable goal using the latest imaging and computational techniques. However, previous approaches have had significant limitations. Temporarily arresting the heart using drugs disrupts the heart’s physiological state, and the use of ultra-high frame-rates for fluorescence image acquisition causes phototoxic cell damage. Real-time triggered imaging, synchronized to a specific phase in the cardiac-cycle, can computationally “freeze” the heart to acquire the minimal number of fluorescence images required for 3D time-lapse imaging. However, until now no solution has been able to maintain phase-lock to the same point in the cardiac cycle for more than about one hour. Our new hybrid optical gating system maintains phase-lock for up to 24 h, acquiring synchronized 3D+time video stacks of the unperturbed heart *in vivo*. This approach has enabled us to observe detailed developmental, structural, cellular and subcellular processes, including live cell division and cell fate tracking, in the embryonic zebrafish heart using transgenic fish lines expressing cell-specific fluorophores. We show that our approach not only provides high spatial and temporal resolution 3D-imaging, but also avoids phototoxic injury, where alternative approaches induce measurable harm. This provides superb cellular and subcellular imaging of the heart while it is beating in its normal physiological state, and opens up new and exciting opportunities for further study in the heart and other moving cellular and subcellular structures *in vivo*.

## 1 Introduction

The ability to image an embryo at a cellular and subcellular level is crucial for understanding the dynamic processes and interactions underpinning development [1–6]. The ideal imaging system would acquire high resolution 3D images of specific organs and structures continuously over relevant time-periods, typically hours or days, without causing tissue damage or interfering with the anatomical or physiological state of the organism. While light sheet microscopy has emerged as a valuable *in vivo* imaging solution to this challenge [7, 8], imaging the complex 3D structure of the beating heart has additional challenges due to its constant cyclic motion at rates of 120-180 beats per minute. This problem can in principle be overcome by using synchronization techniques to acquire 3D images free from motion artefacts [9–11]. However, to date, only some aspects of this problem have been solved at any one time. Imaging of the developing fish and chick heart has been previously reported for: single 3D snapshots at selected intervals[11], 3D images of a single cardiac cycle [12, 13], sampling of a limited number of time points during development [9, 14], and 2D time-lapse video [15]. Temporarily stopping the heart using high dose anaesthetic has also been used to acquire 3D image sets of the heart [16] but such approaches may significantly alter the physiological state of the heart [17]and the wider embryo [18], particularly when performed repeatedly in the same embryo at multiple timepoints.

The challenges of time-lapse 3D cardiac imaging can be understood on three key timescales. Within one cardiac cycle there are rapid changes in shape and size of the heart over less than a second. From one cardiac cycle to the next the heart adopts a highly repeatable sequence of shapes, although there can be subtle changes in rhythm from one beat to the next. However, on longer timescales dramatic structural and functional changes occur at a cellular and whole-organ level, especially during development where the heart undergoes significant morphological changes. It is important to be able to maintain stable imaging over all of these timescales in order to fully understand the biological processes of interest. For example, cell shape changes and cell migration take place over minutes, while cell division and acute inflammatory responses occur over hours. At a whole-heart level, developmental processes such as cardiac morphogenesis take 2-3 days to complete, and injury-associated inflammation also takes several days to resolve.

We have previously demonstrated prospective optical gating in the beating zebrafish heart [10, 11], stroboscopically building up a synchronized 3D z-stack of the heart without motion artefacts (i.e. the imaging is “phase-locked”) such that the heart is at exactly the same phase in the cardiac cycle when every plane is imaged. In contrast to alternative “retrospective” optical gating approaches [9], we only needed to acquire one single fluorescence image per z plane per timepoint. Thus, in the same way that light sheet microscopy eliminates redundant excitation of fluorescence in the spatial domain, the philosophy of our prospective approach is to eliminate redundant excitation in the time domain. This orders-of-magnitude reduction in phototoxic laser dose is a crucial starting point for viable time-lapse imaging. Our previous algorithm assumed quasi-periodic, stereotypical motion from one cardiac cycle to the next and, specifically, in relation to a reference heartbeat. However, on developmental timescales this assumption no longer holds as the heart changes in position, size and shape and correct phase-lock is lost within approximately one hour.

To overcome this limitation we have now developed a new method to maintain day-long, synchronized 3D time-lapse imaging in the beating heart. This has enabled us, for the first time, to track developmental morphogenesis and patterning in the heart with a high degree of temporal and spatial resolution in 3D, and track individual motile cells, including cardiomyocytes and inflammatory cells, associated with the developing and injured heart.

## 2 Results

Our new “hybrid” optical gating algorithm combines prospective optical gating (for synchronized 3D fluorescent stack collection) with retrospective optical gating applied to brightfield images as part of our strategy to maintain phase-lock by adapting to the changing appearance of the heart on developmental timescales. We describe the algorithm as “hybrid” since it combines the strengths of both of these synchronization philosophies. Thus for individual *z* stacks we continue to use our previous prospective gating algorithms [10] to generate the synchronization triggers for image acquisition on every cardiac cycle, based on realtime comparison of brightfield images against an initial reference brightfield image-set representing one cardiac cycle (Figure 1, Supplementary Video 1). However, at regular intervals (typically, after a complete *z* stack has been acquired for the current timepoint) our new algorithm refreshes this reference image-set so that our system can adapt to the changing appearance of the heart (Figure 2) without losing synchronization (Supplementary Figure 1; Supplementary Video 2).

**Figure 1:**
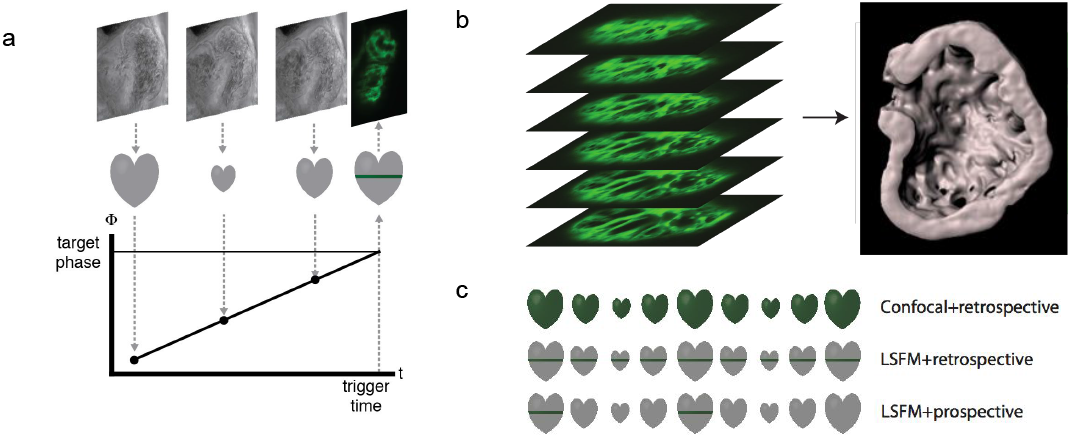
Prospective optical gating minimises specimen exposure to laser light, reducing phototoxic effects. **a,** Brightfield images (greyscale) are acquired and analyzed in realtime to assign a *phase* (temporal position in the cardiac cycle) to each frame. A locally-linear fit to this brightfield phase history over a time window of ~100 ms is used for forward prediction, to anticipate when an electrical signal should be sent to trigger acquisition of one synchronized fluorescence image (green). **b,** As the sample is scanned through the light sheet, a synchronized z stack (green individual images) is generated to yield one 3D timepoint in a timelapse dataset (cutaway rendering of ventricle shown in grey). **c,** To acquire a single 2D image plane in confocal imaging, the entire sample would be illuminated (green shading in diagrams), but in light sheet fluorescence microscopy (LSFM) only the plane of interest is illuminated, reducing phototoxic effects by several orders of magnitude. This same philosophy is applied in the time domain for prospective gating: retrospective heart-synchronization algorithms would require *video imaging* of every single plane, but instead we only trigger image acquisition at the exact desired target phase in the cardiac cycle.

**Figure 2:**
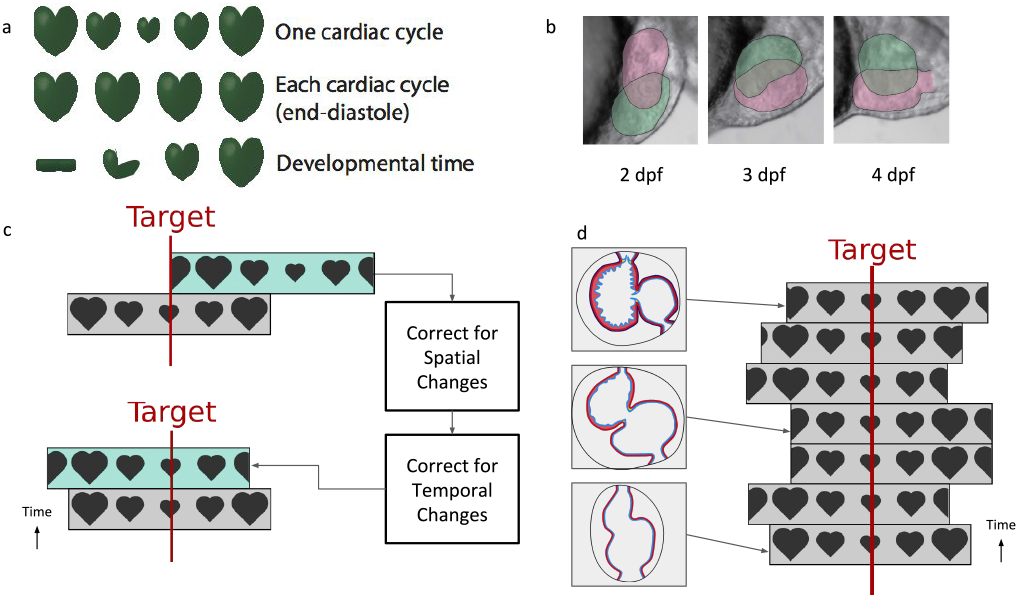
Hybrid optical gating automatically accommodates temporal and spatial changes in heart physiology, permitting stable phase-locked timelapse image capture. **a,** Within each quasi-periodic heartbeat there is substantial cyclical movement (top), and unsynchronized images would be acquired at random, different phases in the cardiac cycle. Prospective optical gating allows synchronized image capture to be triggered every time the heart returns to the same configuration on successive beats (middle). However, over developmental timescales, the heart undergoes structural changes and its appearance is modified (bottom); a long-term synchronisation algorithm must adapt to developmental changes and maintain phase-lock, to permit continuous timelapse imaging over periods longer than ~30 mins. **b,** These developmental changes are evident in the brightfield images used for prospective optical gating. Between 2 and 4 dpf the appearance and structure of the heart (shown here at mid-systole; atrium shaded magenta, ventricle green) changes dramatically; the reference images used for prospective optical gating must therefore be regularly updated in order to maintain correct phase-lock for synchronisation, **c,** Hybrid optical gating offers a solution to this problem. Between acquisition of fluorescence stacks, new reference images are collected (highlighted in upper panel as cyan background) and our automatic temporal alignment algorithm corrects for spatial and temporal changes in order to maintain the same phase-lock. The same target phase of the cardiac cycle selected by the user (in this case, at end-systole) is maintained across all reference images (red line), **d,** This enables us to maintain consistent phase-lock over long timescales (≥24 hours of imaging), despite significant structural and physiological changes (as shown on left).

The key technical advance of our new algorithm is the ability to ensure that the same target phase of the cardiac cycle is maintained over long periods of time (phase-lock), in spite of these regular refreshes of the reference image-set. This is achieved by a suitable cross-correlation between the old and new reference image-sets in the phase domain, taking account of sample movement due to growth (Figure 2c-d; Supplementary Video 3; full algorithm details in Methods). Thus we can optically freeze the heart for however long a specimen can remain biologically viable in the imaging chamber. By ensuring that phototoxicity is minimized and dataset sizes are kept manageable, we are able to perform detailed assessment of structural changes in the developing heart, and visualise cellular and subcellular behaviour, near to continuously, over the course of hours and days.

### 2.1 Hybrid Optical Gating Allows the Capture of Developmental Changes in the Living Zebrafish Heart Despite Changes in Shape, Size and Rhythm

The zebrafish heart tube forms and begins contracting rhythmically by 24 hpf. This linear tube elongates asymmetrically and by 36 hpf the future ventricle migrates ventrally in relation to the atrium, a process termed cardiac looping [19, 20]. By 48 hpf, the looped zebrafish heart appears S-shaped and undergoes more subtle changes until both chambers are morphologically distinguishable [19, 21]. Using our hybrid optical gating software, we have observed these changes in the heart, and associated developmental changes in the nearby vasculature from 48 hpf-72 hpf. By acquiring a z-stack every 5 mins, we observed the pinching of the atrioventricular canal region, followed by expansion of the outer curvature of the atrium and finally the ventricle (Figure 3a; Supplementary Videos 4-5). This enlargement is associated with a 300% increase in endocardial chamber volume between 48 hpf and 72 hpf. To our knowledge, this is the first reported 3D time-lapse movie detailing cardiac morphogenesis of a live beating zebrafish embryonic heart.

**Figure 3:**
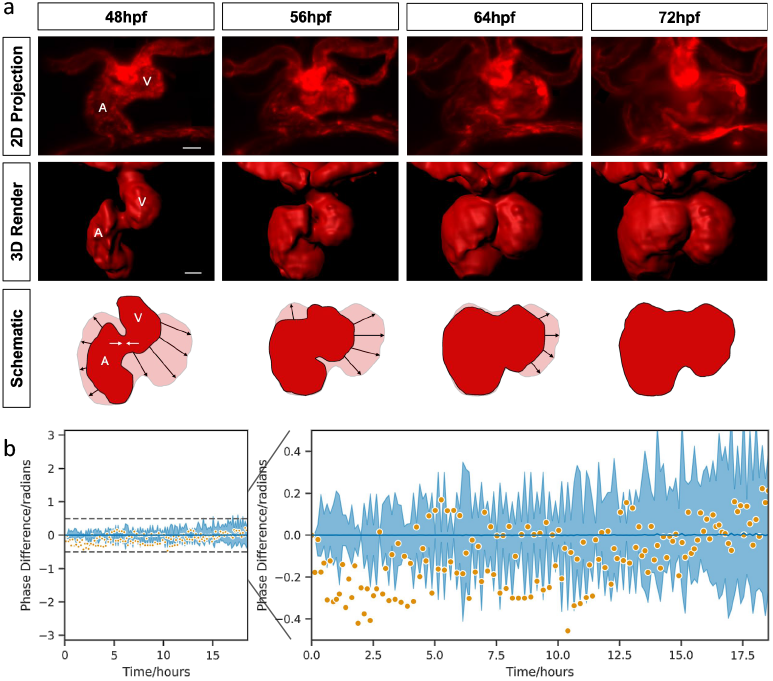
Hybrid optical gating allows phase-locked, long-term, time-lapse imaging and quantification of developmental cardiac morphogenesis. **a,** Phase-locked, live time lapse imaging of cardiac developmental changes (48-72 hpf, at 300s intervals) showing endothelium lining blood vessels and heart chambers (red - transgene *flk1:mCherry*). Selected timepoints shown as maximum intensity projections of z-stacks (top; from Supplementary Video 4); 3D rendering of heart chambers (middle; from Supplementary Video 5); schematic representations of the atrium and ventricle at each timepoint, with final morphological conformation of heart for reference in pale red (bottom). Black arrows represent direction of chamber expansion and white arrows represent atrioventricular pinching, V = ventricle, A = atrium. Scale bars: 30*μ*m. **b,** Hybrid optical gating algorithm (orange) phase-lock performance compared against mean human judgements of best-matching frames (blue; n = 3 per timepoint, line represents mean and shaded area represents the standard deviation). Throughout an 18 hour dataset (during the dramatic changes of cardiac looping) there is only small variability between automated and human results (left - full 27ĩ range). Zooming in (right) we can see that, for this dataset, some drift in phase occurs over time but within ±0.5 radians (approximately 3 brightfield frames or 40 ms). Apparent variations in the automated synchronisation algorithm are similar to the spread observed in the manual human assessment.

To explore the accuracy of our hybrid optical gating software we compared the software-based frame selection of a specific phase in the cardiac cycle with a human-based manual selection during a 24 hour time-lapse experiment (Figure 3b). We found strong agreement between the target frame chosen manually by five human experts (searching for a fixed target frame in the cardiac cycle - see Methods) and that chosen by our fully automated software algorithms: our algorithms maintain a locked target phase with similar variance to human experts (*σ*^2^ = 0.022 vs 0.037 radians for human experts) despite significant developmental changes in cardiac shape and size. We note that neither the human experts nor fully automated algorithms can be considered a “ground truth”, and often human experts disagreed among themselves in identifying the best-matching target frame, by several frames.

### 2.2 Hybrid Optical Gating Allows 3D Time-lapse Imaging at a Rate Fast Enough to Track and Assess Inflammatory Cell Behaviour

We have also studied the migration of neutrophils and macrophages onto and into the beating heart following targeted laser-injury to the apex region of the ventricle [23]. Our phase-lock imaging capabilities permit high spatial resolution while imaging inflammatory cells in contact with the heart. We observed minute-by-minute changes in inflammatory cell shape, size, velocity and behaviour as they migrate to and from the injury site on the beating heart (Figure 4a-c; Supplementary Video 6, 7). From datasets such as these we can accurately quantify inflammatory cell behaviour in 3D, tracking individual neutrophils migrating to and from the injury site (Figure 4d, Supplementary Figure 2b).

**Figure 4:**
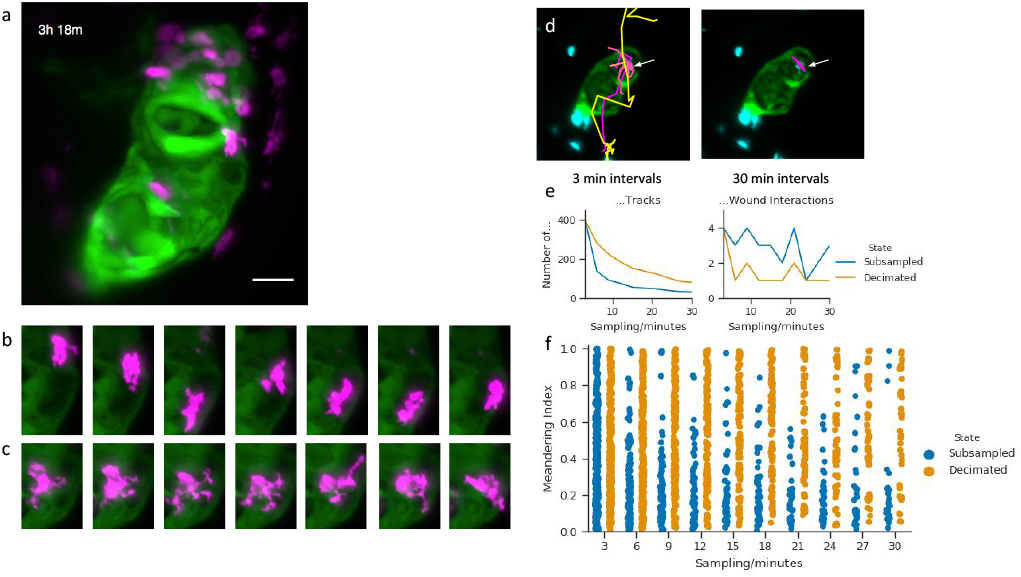
Hybrid optical gating allows phase-locked time-lapse imaging at the rate required for investigating immune cell activity during injury response. **a,** Maximum intensity projection of z-stack showing macrophages (magenta - transgene *mpeg:mCherry*) at a site of laser-injury on the upper pole of the heart ventricle (green - transgene *myl7:GFP*). Image is a still frame from Supplementary Video 6. Injury was induced at 74 hpf, with imaging commencing at approx 83.5 hpf. Scale bar: 30*μ*m. **b,c,** Selected maximum intensity projections highlighting complex shape changes during migration in two individual macrophages (displayed at 6 minute and 3 minute intervals, respectively), **d,** Maximum intensity projection of z-stack showing a neutrophil (cyan - transgene *mpx*:mCherry) at the injury site on the apex of the ventricle (white arrow) (green - transgene *myl7:GFP*). Image is a still frame from Supplementary Video 7. Injury was induced at 72 hpf, with imaging commencing at approx 74 hpf. The high temporal acquisition rate permits track analysis (coloured lines) of rapidly migrating neutrophils (cyan - transgene *mpx:mCherry*) migrating to and from the laser-injury site (white arrow). At 3 minute intervals (left), a rate only feasible with our hybrid optical gating, we can clearly track two neutrophils that interact at the wound site; however, at 30 minute intervals (right), as commonly used in other longitudinal imaging studies, only one neutrophil is seen to interact and the captured track misses the detailed migratory behaviour of that neutrophil, **e,** Poorer sampling intervals significantly degrade the accuracy of track recovery (left) and the number of detected wound region interactions (right). These plots examine tracks recovered from temporally degraded data: either the timelapse image data was temporally subsampled before track analysis (blue), or tracks computed from 3 minute interval image data were subsequently temporally decimated (orange). Reasons for considering these two degradation modes are explained in Online Methods, **f,** Track errors at longer sampling intervals cause quantification errors when trying to assess neutrophil behaviour. Here we show errors in the meandering index [22] computed for tracks recovered from both subsampled image data (blue) and decimated tracks (orange); the meandering index quantifies the tortuosity of a neutrophil track.

The short interval of 150s between each complete 3D stack, made possible by our hybrid imaging system, allows detailed tracking of highly motile neutrophils (Figure 4e-f) without risk of photo-toxicity or bleaching. Figure 4f illustrates how neutrophil migratory behavioural information could be readily lost by using longer time intervals between z-stacks as shown by changes in *meandering index* [22] as the 3D image acquisition rate falls. The meandering index quantifies the tortuosity of a neutrophil track, i.e. whether the motion of a neutrophil is highly directed (which could for example represent fast movement through an adjacent blood vessel) or more tortuous (a potential signature of cell-cell interaction at the wound site). For the purpose of this demonstration we draw the reader’s attention to two features of this plot: 1) As the sampling interval is increased, both curves show considerable changes in the number of tracks and value of meandering index; this is mostly due to very fast neutrophils having transited the field of view between frames. In this situation a large sampling interval is leading to an unrecoverable loss of information. 2) At longer sampling intervals the two curves differ; this is mostly due to erroneous automatic track recovery in longer sampling intervals of subsampled 3D videos compared to the “ground truth” of decimated tracks. Here a large sampling interval prevents or hinders automatic image analysis, making quantification difficult and possibly inaccurate.

The inflammatory cell response following heart injury, involving both neutrophils and macrophages, typically takes more than 24 hours to fully resolve. Our hybrid optical gating approach is essential to provide the spatial and temporal resolution needed to understand this dynamic process over this time course.

### 2.3 Hybrid Optical Gating Reduces Photo-injury and Photobleaching Compared to Retrospective Optical Gating Approaches

A key advantage of our hybrid optical gating approach is that it minimises the number of fluorescence images acquired, and hence minimises the deleterious effects of laser exposure on the tissue (see [24, 25] for recent discussions). Light sheet fluorescence microscopy is well known for its ability to reduce photobleaching and photodamage by limiting fluorescence excitation to only the spatial plane(s) of interest [26]; hybrid optical gating further limits fluorescence excitation to only the times (i.e. cardiac phase) of interest. In addition, this reduction in laser dose minimises the risk of photodamage-induced changes in heart rate and rhythm, which would present additional challenges for high-quality phase-locking.

To quantify the physiological impact of established retrospective gating strategies in comparison to our hybrid optical gating, we used heart rate as a measure of photo-injury using these two different imaging protocols (see Methods). For embryos exposed to two hours of retrospectively-gated imaging at 5 minute intervals, we observed an initial increase followed by a significant reduction in heart rate. We speculate that this is due to an initial heating, causing an increase in heart rate, followed by a patho-physiological response, resulting in bradycardia, due to photodamage and phototoxicity to the heart. We also observed changes in heart rhythm (n=5 of 6 fish) and reduction in ventricle ejection fraction (Supplementary Video 8). In contrast, time-lapse imaging using our hybrid optical gating at 5 minute intervals caused no significant change in heart rate over a similar time period (Figure 5a).

**Figure 5:**
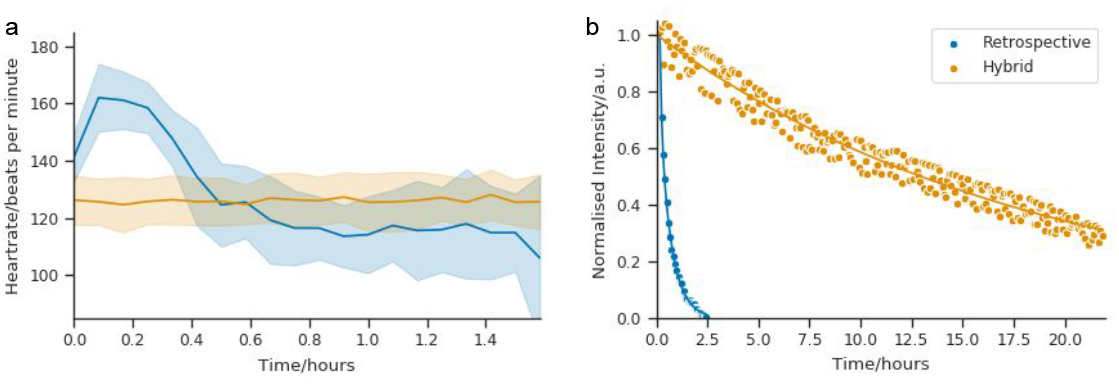
Hybrid optical gating protocol causes less cardiac photo-injury and photo-bleaching than retrospective gating protocols. **a,** Acquisition of multiple fluorescent z-stacks in an embryonic heart (transgene *myl7:gfp*; 3 dpf) using a retrospective optical gating protocol (blue; mean (line) with standard deviation (band)) causes an increase, followed by a fall, in heart rate over a 1.6 hour imaging period (n=6 fish). Meanwhile 1.6 hours of continuous image acquisition using our hybrid optically-gated protocol (orange; mean (line) with standard deviation (band)) with stacks acquired every 5 min, causes no change in heart rate (n=5 fish). Plots for individual fish can be seen in Supplementary Figure 3. **b,** When exposed to light levels required for retrospective gating (using typical parameters reported in the literature, and again acquiring z stacks at 5 minute intervals), the fluorescence signal from the highly bleachable membrane-specific mKate fluorophore (transgene *myl7:mKate-CAAX*) is halved after only 3 z-stacks (blue; mean (line) and individual frames (points)). In contrast, hybrid optical gating achieves 150 z-stacks before showing an equivalent fall in fluorescence intensity (orange; mean (line) and individual frames (points)), thus permitting sustained timelapse imaging. See also Supplementary Video 9. Data shows one representative fish for each condition.

The reduced laser dose with hybrid optical gating also has an important impact on the rate of photobleaching. We demonstrated this by acquiring 3D time-lapse fluorescence images using two alternative imaging protocols (Figure 5b, Supplementary Video 9). The first protocol, adopting a retrospective optical gating approach, causes rapid and marked bleaching of mKate emission from cardiomyocyte cell membranes and the signal soon becomes indistinguishable from background. In contrast, our hybrid optical gating protocol causes only mild and gradual photobleaching, and can continue to capture 3D images for over 24 hours. Direct reduction in bleaching rate is due to a combination of the stroboscopic nature of imaging [27] and the reduced total laser dose [28]. Additionally, a reduction in photo-injury in the tissue may minimise the oxidative rate of fluorescent molecules, as previously seen in *in vitro* samples [29], thus indirectly reducing photobleaching still further.

### 2.4 Hybrid Optical Gating allows live observation of cardiomyocyte cell division and provides novel insights into myocardial trabeculation

To demonstrate the advantages of our new platform for addressing complex biological questions, we studied the process of cardiac trabeculation through time-lapse imaging of cardiomyocyte migration and cell division events in the beating heart (Figure 6a,b; Supplementary Video 10). Trabeculation is the developmental process whereby multicellular luminal projections of cardiomyocytes are formed. Studying trabeculation in vivo has thus far been limited by imaging techniques that risk perturbing the heart if applied at the temporal resolution and duration required to understand this 3D cellular patterning [5, 30]. Using our approach, we were able to monitor trabeculation at the tissue level in the same embryonic heart during three consecutive days. A luminal view of the surface-rendered myocardium shows, in detail, the development of trabecular patterns as the heart matures (Supplementary Figure 4a).

**Figure 6:**
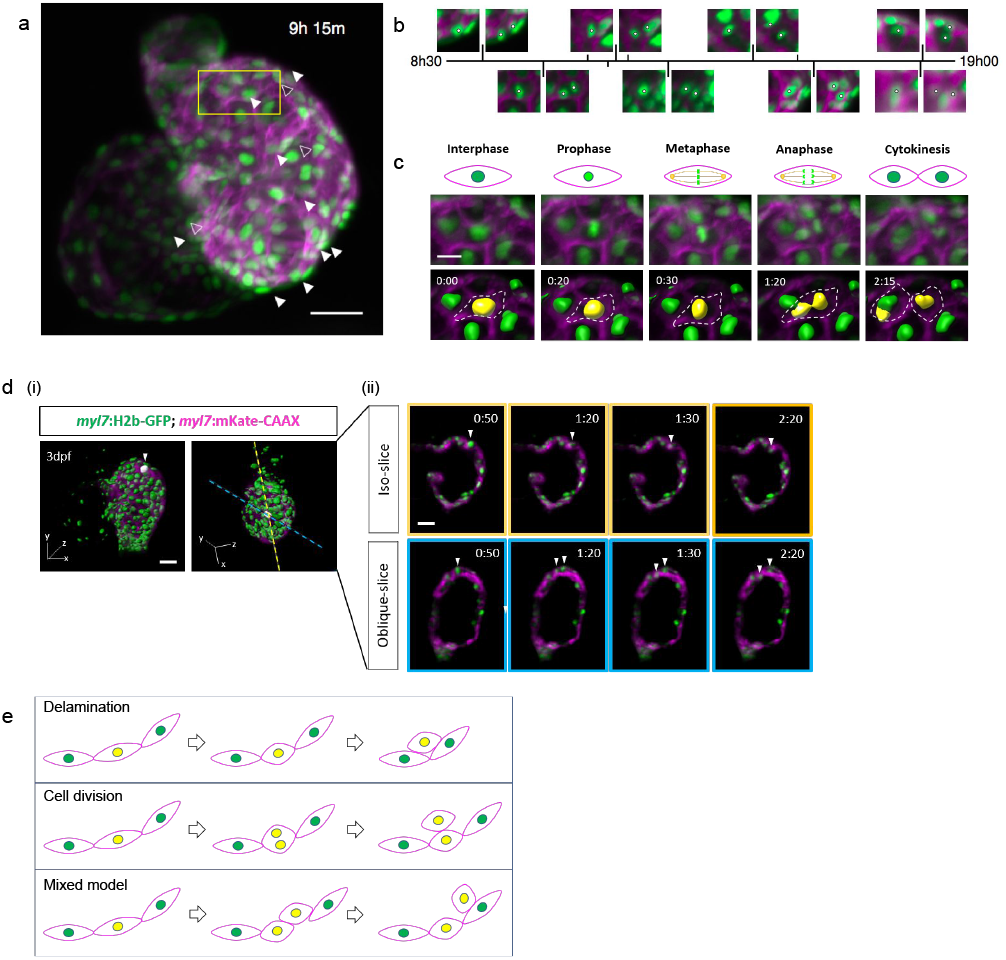
3D time-lapse imaging accurately maps cardiomyocyte migration and cell division during myocardial trabeculation. **a,** Representative MIP from a 24 hour timelapse of a 72-96hpf embryo (Supplementary Video 10). Image shows cardiomyocyte nuclei (green - transgene *myl7:h2b-gfp*) and cardiomyocyte cell membranes (magenta - transgene *myl7:mKate-CAAX*), with strongest signal expressed in the ventricle. Nuclei of cardiomyocytes which will later divide are indicated by white arrowheads (open-arrowheads for less-obvious divisions near the posterior wall of the heart). Region of interest for c is indicated with yellow box. **b,** detail from timelapse: pairs of images plotted on a timeline, illustrating selected cardiomyocytes shortly before (left) and after (right) cell division; nuclei indicated with white circles, **c,** A selected cardiomyocyte division clearly exhibits key stages of mitosis. Mitotic stages (schematic: top row) are visible at selected timepoints from the timelapse sequence (MIPs: middle row), and the nuclear volume changes and karyokinesis typical of cell division can be seen (dividing nucleus surface-rendered in yellow: bottom row), **d,** (i) Whole embryonic heart ventricle with nuclei surface-rendered (green) and superimposed onto a MIP of cell membranes. Nucleus of cardiomyocyte destined to divide (left) is shaded white and indicated by an arrowhead; original plane of slice acquisition (yellow dashed line) and plane of nuclear division (blue dashed line) are also indicated, (ii) the dividing cardiomyocyte nucleus is shown in the acquisition plane ‘isoslice’ view (top row) and in the plane of division ‘oblique slice’ view (bottom row). In the latter the white arrowheads highlight the observation of two daughter cardiomyocyte nuclei, which would have been missed in the isoslice view, **e,** Schematic illustrating three models of cardiac trabecular initiation: delamination model (top), cell division model (middle) and mixed model (bottom). Delaminating cardiomyocytes are indicated by a yellow nucleus. Scale bars: 30*μ*m; 10*μ*m for **c.**

Two principal mechanisms have been proposed for initiation of trabeculation [30]. The first is “delamination”, where cardiomyocytes are physically squeezed-out luminally from the cortical myocardium layer without a cell division event. The second is “orientated cell division”, where cortical cardiomyocytes divide perpendicular to the ventricular myocardium. Currently, delamination is suggested to be the sole mechanism of trabeculation initiation in zebrafish [30]. Using our imaging system we were able to assess whether orientated cell division could also contribute to trabeculation, using the *Tg(myl7:h2b-GFP; myl7:mKate-CAAX*) zebrafish line, which fluorescently reports both cardiomyocyte chromatin and cell membrane. We acquired phase-locked time-lapse z stacks at 10-minute intervals from 72-96 hpf, allowing us to observe the archetypal stages of mitotic cell division at both the sub-cellular and cellular level (Figure 6c). During cardiomyocyte cell division chromatin can be seen to condense, align on the metaphase plate and is then pulled to opposite poles to form two daughter cells which finally become separated by plasma membrane.

We observed migratory patterns of ventricular cardiomyocytes and cell division events occurring both parallel and perpendicular to the abluminal-lumen axis (Supplementary Figure 4b). Interestingly, tracking the migration of one such cardiomyocyte in the original plane of acquisition (Figure 6d(i)) initially indicated that it delaminates from the cortical layer without dividing (iso-slice, Figure 6d(ii)). We undertook further analysis by re-slicing the full z-stack along additional planes (Figure 6d(i) - blue dashed line) which revealed that this cardiomyocyte actually undergoes cell division, parallel to the abluminal-lumen axis, whilst also delaminating perpendicularly (oblique-slice, Figure 6d(ii)) creating a trabecular projection into the lumen. These observations potentially suggest a “mixed model” for trabeculae formation in the ventricle, whereby cardiomyocytes divide *and* then migrate luminally (Figure 6e). The frequency and extent of such cardiomyocyte behaviour clearly requires further study but it highlights the critical importance of examining myocardial growth patterning in an imaging system capable of high resolution 4D imaging.

## 3 Discussion

The key advance that distinguishes our approach from retrospective synchronization strategies is that we actively trigger image acquisition based on decisions made in real-time, thus eliminating all unnecessary acquisitions of fluorescence images [31]. Retrospective strategies typically require several hundred fluorescence images at every z-plane [13], but if the aim is to “freeze” the heart at a desired phase in its cycle then only one of these images is finally selected. All other images represent unnecessary exposure of the sample to laser excitation light, thus accelerating harmful phototoxicity and photobleaching. Light sheet microscopy reduces the damaging laser exposure by orders of magnitude compared to confocal imaging, but this benefit is lost when large numbers of fluorescence images must be acquired in each plane prior to retrospective synchronization. Similarly, pharmacological arrest of the heart using high dose anaesthetic permits artefact-free imaging, but at the cost of exposure of the heart and the whole organism to repeated exposure to anaesthetic agent.

Retrospective strategies do have the advantage of allowing freedom to make *post hoc* decisions to reconstruct images from all phases of the cardiac cycle, which is advantageous for studies of cardiac dynamics [32]. However, for developmental and cell migration studies, as presented here, only images at a single phase need be acquired. We note that if additional phases in the cardiac cycle are required, for example for ejection fraction measurement, blood flow mapping [33], or studies of heart wall dynamics, then our approach is readily adapted to generate images at multiple precise phases throughout the cardiac cycle. Recent advances in fast volume imaging using custom-built adaptations of light sheet microscopes [34–36] may soon permit direct, synchronization-free “snapshot” 3D imaging of the heart. Even as these techniques mature, for time-lapse imaging our real-time triggering capability will still remain essential for triggering repeated time-lapse 3D image acquisitions at a *fixed* phase in the cardiac cycle.

A further important advantage of our method is that it substantially reduces data file size and thus facilitates data handling, particularly for time-lapse imaging. For example, if a retrospective gating approach had been used for the experiments in Figure 3 and Supplementary Video 4, the raw data would have occupied at least 16 TB (assuming 200 video frames per slice [13]), and would have required many hours of post-processing time. In contrast, our approach requires only 80 GB of storage space (24 z-stacks per hour for 24 hours), thus significantly reducing the challenges of storage and processing of large datasets typically acquired in time-lapse 3D imaging. These modestly-sized z-stack files can also be accessed and reviewed by the user through the course of the experiment, paving the way for future developments such as user-initiated or automated responses to unexpected or atypical biological observations during imaging experiments.

We have demonstrated that our optical-gating system can be used to clarify complex biological events such as the cellular processes underlying myocardial trabeculation. Two previous studies have used genetic tools to stochastically label cardiomyocytes with different fluorophores prior to trabeculation [30, 37]. These studies reported that adjacent cortical-trabecular cardiomyocytes rarely share the same fluorophore, implying that orientated cell division does not contribute to trabeculation. However, these studies relied on analysis of single optical and histological slices of the heart and so may have missed daughter cells that have migrated away from each other out of plane. Our own observations have shown that in a number of instances cardiomyocyte cell division was followed by immediate perpendicular migration of a daughter cardiomyocyte. Furthermore, coupling delamination to proliferation is a plausible strategy for trabeculation. Not only would this maintain cortical cardiomyocyte number, the disassembly of sarcomeres (known to occur prior to proliferation) might also liberate the cell for delamination. Our findings suggest that such events could have been missed in studies where time-lapse imaging was acquired only in a single plane, potentially underestimating the contribution of dividing cortical cardiomyocytes to pioneer trabecular cardiomyocytes [5, 6, 30, 37]. This insight was only possible due to the high spatial and temporal resolution of our optically-gated imaging system.

The inflammatory cell behaviour we observed following heart laser injury also highlights the value of our method for 3D time-lapse imaging. The ability to track individual, rapidly moving inflammatory cells on the injured heart provided novel insights into neutrophil migratory behaviour on the beating heart. There is a clear potential to study pharmacological and genetic manipulation of these aspects of neutrophil migratory behaviour at the cardiac injury site. Similarly, changes in macrophage cell morphology were observed using our system, highlighting the potential to study morphological changes in heart-associated macrophages *in vivo*. Linking these shape changes with specific molecular phenotypes could provide new insights into macrophage subsets in the injured heart.

Our multi-modality approach, using brightfield images to trigger synchronized acquisition of the fluorescence image, is not only useful for real-time synchronization but also separates the conflicting requirements for fluorescence imaging (tissue-specific labelling, which may be highly sparse; z-scanning; z-sectioning while minimising photodamage) and heart-synchronization (phase-diverse periodic signal; invariance during z-scan), offering additional flexibility to optimize future microscope designs for cardiac imaging. This is illustrated by our ability to conduct high-precision flow mapping by fusing information across multiple cardiac cycles in a statistically-rigorous manner [33]. In these experiments the periodic nature of the flow is less directly discernible from the raw fluorescence data, but easily obtained from the accompanying brightfield channel. Using a brightfield synchronization source also has other potential uses such as fusing and phase-registering multiple fluorescence channels acquired sequentially in the case of a microscope with a limited number of fluorescence imaging ports for example or acquiring synchronized images in multi-acquisition modalities such as structured illumination microscopy.

Our hybrid gating approach is equally applicable to embryos of other species such as fly, chick or mouse. Transferring this technology to another microscope system, such as 2-photon or confocal microscopes, is also possible and would simply require access to a brightfield image channel with the ability to trigger image acquisition. Our synchronization software can run independently of the existing microscope control software, avoiding any need for complex integration at software level. Thus, with a small degree of cooperation from manufacturers over hardware integration, our system can be integrated with commercial two-photon systems (demonstrated in Supplementary Figure 5) and light sheet microscopes, as well as custom-built microscope platforms, to equip them with real-time synchronization capabilities.

In conclusion, we have shown that our hybrid optically-gated imaging system can provide high spatial and temporal resolution time-lapse 3D images of the beating heart, as demonstrated here for embryonic zebrafish. 3D images can be readily obtained throughout 24 hour time-lapse studies, allowing us to observe cardiac morphogenesis, inflammatory cell migration and cardiomyocyte cell division. This approach has the potential to provide further novel insights into morphological, cellular and subcellular processes in the beating embryonic heart in the zebrafish and other species.

## 4 Acknowledgements

This work was funded by the British Heart Foundation (BHF) (NH/14/2/31074), with additional support from the BHF (RE/08/001), EPSRC (EP/M028135/1) and the Royal Society (RG130249). Individual support: FB & CB (BHF CoRE award RE/13/3/30183), AB (Medical Research Scotland project ID 3376798), CJN (EPSRC Doctoral Prize Research Fellowship EP/N509668/1), JMT (Royal Society of Edinburgh Sabbatical Research Grant RSE58915). We thank Dr Thai Truong (University of Southern California) for proposing the side-view geometry used for brightfield imaging.

## 5 Author contributions

JMT and CJN developed the synchronization algorithms, software, electronics and optics. CJN, FB, AB and CB performed the experiments and quantitative analysis. CJN, FB, AB and CB conceived individual experiments and developed the protocols. JMT, MAD and JM conceived the study and supervised the research. The manuscript was written with input from all authors.

## 6 Competing financial interests

The authors declare no competing financial interests.

## Methods

### 1 Fish husbandry and embryo preparation for imaging

All experiments were performed in accordance with the Animals (Scientific Procedures) Act 1986 in a UK Home Office-approved establishment. Zebrafish husbandry, embryo collection and maintenance were performed according to accepted standard operating procedures [38]. Briefly, embryos were maintained in embryo medium (1 ×conditioned water (CW) containing 0.1% (w/v) methylene blue) at 28.5 °C on a 14 h light/10 h dark cycle and all experimental procedures were performed at room temperature (23 °C). Transgenic zebrafish lines used for imaging are as follows; Tg(*myl7:eGFP*^twu26^) [39], Tg(*kdrl:mCherry*^ci5^) [40], Tg(*mpx:mCherry*^uwm7^) [41], Tg(mpeg1:mCherry) [42], Tg(*myl7:h2b-GFP*^zf52^) [13], Tg(*myl7:mKate-CAAX*^sd11^) [43]. Adults were day-crossed as appropriate to yield desired combinations of transgenes in embryos. Embryos were treated with phenylthiourea (Fisher Scientific) at 7 hpf to inhibit pigment formation and enhance image clarity. Embryos were embedded in 0.5% low-melting-point agarose inside FEP tubes (Adtech Polymer Engineering) to prevent drift whilst still allowing growth during long term imaging. Fish were used only once for a timelapse imaging experiment, and any repeats shown come from distinct individuals.

### 2 Light sheet microscope set-up

Our custom-built selective plane illumination microscope (SPIM) is optimized for simultaneous multi-channel light sheet fluorescence imaging of zebrafish (Supplementary Figure 6). The detection arm is based around a Nikon Plan Fluorite 16x/0.8 N.A. objective lens with ultra-flat dichroic mirrors (Chroma T495lpxr-UF2, T550lpxr-UF2, T700spxr-UF2) separating red/green/blue fluorescence channels, and a near-IR, non-visible brightfield channel (which is not routinely used - see below). Fluorescence images are acquired using QIClick CCD cameras (QImaging) and fluorescence emission filters (Thorlabs MF479-40, MF525-39, MF630-69). Brightfield images are normally acquired by imaging through the light sheet launch objective (for reasons discussed in Supplementary Note 1), with a 650nm shortpass dichroic mirror (Edmund Optics) serving to pick off the collected brightfield light, which is imaged onto a Prosilica GS650 CCD camera (Allied Vision).

Laser excitation is at 488 nm and 561 nm, using a Versalase laser system (Vortran) with single-mode fiber delivery. Brightfield illumination is via a light emitting diode at 750 nm (near infrared, non-visible). The light sheet (measured full-width-at-half-maximum 2.4 *μ*m) is formed using a cylindrical lens and a 10x/0.3NA objective lens (Nikon). Shadow effects are minimized using an mSPIM configuration [44] using a 4 KHz resonant scan mirror (SC-30, Electro-optical Products Corporation). Laser pulsing and camera triggering is coordinated using a custom-built electronic system based around a StartKit microcontroller board (XMOS; software and hardware details in Supplementary Note 2) which also serves to define a universal timebase for our synchronization analysis. The sample is positioned and scanned using a combination of manual micrometer stages and motor-driven stages (M-111.1DG, Physik Instrumente).

Imaging parameters used for all experiments can be found in Supplementary Table 3.

### 3 Real-time, prospective optical gating

Real-time, prospective optical gating [10, 11] is used to trigger capture of fluorescence images at a user-selected target phase in the heart’s natural cycle. This approach allows us to computationally freeze the heart without any invasive approaches such as pharmaceuticals or pacing. Real-time, prospective optical gating relies on information captured in the brightfield channel (for which even continuous illumination is not sufficiently intense to cause harm to the specimen). First, a set of reference brightfield images is collected, lasting precisely one natural heart cycle. Subsequent brightfield images are correlated against this to assign them a phase. A target phase in the heartbeat is selected by the user, and all subsequent fluorescence images will be triggered at this target phase in the heartbeat.

For each brightfield image *I_t_* (composed of pixels *I_t_*(*x,y*)) received at the current time *t*, the following steps [10, 11] are taken to phase-match the heart images and forward-predict the next fluorescence trigger time:

1. The new brightfield image is windowed to correct for any in-plane (xy) sample motion Δ*x*, Δ*y*, as calculated from the previous-received frame (see below).
2. The windowed brightfield image is compared against the one-heartbeat reference image sequence *R_ϕ_* to identify the current phase *ϕ_t_* of the heart using a sum of absolute differences metric (this metric was chosen to maximise realtime processing speed):

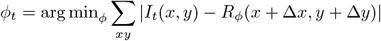
3. The phase *ϕ_t_* is refined to sub-frame precision using V-fitting as described in [10].
4. To accommodate unavoidable latencies in the system, linear forward-prediction of the phase evolution is used to calculate the required triggering time for the next acquisition, i.e. the precise time at which the heart will next be at the desired target phase of its heartbeat.
5. A decision is made whether to commit to the calculated trigger time or to await a refined prediction from the next incoming brightfield image. This decision is made based upon the known processing latency (~12ms from brightfield image exposure to programming of a trigger).
6. The measure of uniform drift Δ*x*, Δ*y* in the *xy* plane of the brightfield images (as used in step 1) is updated as described in [11], ready for the next incoming brightfield image.

If the system decides to send a trigger, the timing details are transmitted to the timing controller over a USB/serial link. Once the timing controller has been informed of the required trigger time, it will electrically trigger the next, synchronised fluorescence image. Following this, the sample is moved to the next z-position in preparation for the next synchronised trigger.

When assigning a phase to the current brightfield image, by comparing it to the reference image-set, nonperiodic variations (such as the locations of blood cells) may result in a situation where the best match is with the very first (or last) frame in the reference image-set. This may lead to erroneous phase values being computed. We mitigate against this by “padding” our reference image-set with two extra frames at either end; these extra frames are not used for the initial minimum-finding, but are available in the case where the initial minimum lies at an extremum of the reference image-set.

Forward prediction is achieved by extrapolating a linear fit of the recent measured phase history as a function of time [10]. In principle the number of datapoints used for this linear fit will affect the accuracy of the synchronization, and we hypothesize that limiting the fit to phase values obtained from the current heart contraction only may result in less variability – however in practice our latencies are low enough that neither the precision nor accuracy of the synchronization are seriously affected by the use of different fitting strategies.

### 4 Hybrid optical gating for long-term, time lapse imaging

Our described prospective optical gating approach is based around image similarity metrics (step 2 above). This has the benefit of not making any assumptions about the shape, orientation or appearance of the heart (in contrast to methods in other fields that are based on, for example, edge detection and tracking). However, gradual development or growth of the embryo will lead to changes in the appearance of these images - thus compromising the effectiveness of the synchronisation over time and limiting the utility of a given reference heartbeat to a maximum of around 60 min of synchronisation (Supplementary Figure 1, Supplementary Video 2). To extend this and allow ‘phase-locked’ time-lapse imaging over tens of hours we have used concepts from post-acquisition, retrospective optical gating [12] to update the brightfield reference heartbeats at regular intervals [45] (Supplementary Video 3). Updating the reference heartbeat allows the system to cope with changes in heart size and morphology that occur throughout development and growth.

First, a new set of brightfield reference images must be selected as previously described [10]. We then determine the equivalent target frame in this new reference heartbeat that matches the original, user-defined, target heart phase. This is done by cross-correlating between the original and new reference heartbeat image image-sets as follows:

1. Each reference heartbeat (*R_u,ϕ_*(*x,y*) for timepoint *u*) is re-sampled 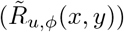 to contain a fixed, integer number of frames.
2. The relative phase shift Δ*ϕ_ab_* is computed between this new reference heartbeat and each of several (typically three) recent past reference heartbeats. This shift is computed by minimising a least-squares criterion representing the similarity between mutually-shifted image-set, as follows:

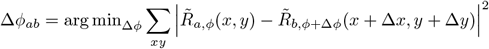 In fact this can be shown to be equivalent to the following criterion, based on crosscorrelation along the phase axis, which we compute in Fourier space for speed:

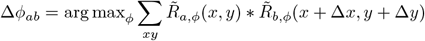
3. Each Δ*ϕ_ab_* is refined to sub-frame precision using V-fitting as described in [10].
4. All the computed Δ*ϕ_ab_* are incorporated, along with relative phase shifts between historical pairs of past reference frames, into a weighted, linear least-squares regression to determine a global phase shift that corrects for small and accumulated errors in the determination of relative phase shifts between individual pairs of reference heartbeats.

We note the importance of accounting for motion (Δ*x*, Δ*y*) of the sample in the xy plane of the brightfield channel. This may be due to growth or movement of the fish within the sample holder. Without correction for this drift, the cross correlation will lose effectiveness due to a reduction in pixel-wise similarity. This will lead to errors in the relative phase shift that would accumulate in the global phase shift over time. Since we already track the drift during the real-time phase matching process (see above, and as introduced in [11]), we are easily able to correct for this drift when comparing pairs of reference heartbeats.

This hybrid of prospective optical gating for real-time phase matching, and retrospective optical gating for long-term phase-locking, allows us to capture 3D time-lapse data at a fixed, specific target phase over extended time periods of 24 h or more.

### 5 Manual Comparison of Long Term Phase-Locking

To verify that hybrid optical gating maintains a consistent phase-lock throughout longterm time-lapse imaging, we asked experts in zebrafish heart imaging (n = 5) to manually identify the same phase in a heartbeat across all the reference heartbeats used to capture the heart looping dataset in Figure 3. Stacks were captured every 150 s using the real-time, prospective optical gating technique described above; automated long-term updating of the reference heartbeat, also described above, was carried out after every third stack. Experts were provided with reference image-sets in a random order, and the sequences began at different phases of the heartbeat. Each individual reference heartbeat was assigned a target frame by n = 3 experts. These manual targets, *T_h_*, were then used to compare the fully automated, hybrid optical gating results. Figure 3 shows these manual targets on a stack-by-stack basis (blue area) as a difference from the human mean (blue line). The fully automated, hybrid optical gating results are also shown on this scale (orange points). In this comparison, neither the human nor the automated updates should be considered as the “gold standard”, but the close agreement between the two is further evidence that a consistent phase-lock has been maintained throughout the 18 h experiment.

### 6 Assessment of photo-injury during imaging

To assess the effect of retrospective optical gating and hybrid optical gating on the physiology of the fish we compared the heart rate of fish in two different scenarios:

1. for fish undergoing two hours of fully automated, hybrid optically-gated time-lapse acquisition (gated stacks acquired every 5 minutes), and
2. for fish undergoing two hours of retrospective optically-gated time-lapse acquisition, acquired in a similar manner to that described in [13]. We used our existing light sheet set-up, but with the laser turned on continuously, to provide illumination equivalent to the acquisition of 600 video frames (each 2.5ms exposure) per z plane (full details in Supplementary Table 1).

Heart rate was calculated every 5 minutes using the period-determination codes described in [11]. Any heart rate determined that varied by more than ±25 % from the previous heart rate value was discounted and a new period determined, which always varied by less than this amount. Such events occurred infrequently and were due to changes in heart rhythm causing erroneous period determination; 8 of these events occurred, all during the retrospective optical gating protocol.

For each protocol (hybrid optical gating and retrospective optical gating) fish were randomly selected from a full clutch of eggs. All fish were given 30 min to acclimatise, from which a resting heart rate was determined. Fish were then imaged for two hours and the heart rate measured at the start of each new stack.

For the hybrid optical gating protocol, fish were mounted in the microscope and, after acclimatisation, were exposed to pulsed laser illumination to accompany triggered (synchronized) fluorescence image acquisition. For retrospective gating experiments, fish were mounted in the microscope and, after acclimatisation, were exposed to constant laser illumination as required for retrospective optical gating (see above). In all cases the fish were also exposed to the infrared illumination needed for brightfield imaging (required for heart rate determination as well as for synchronization analysis).

We note that control fish exposed to no laser light at all show no change in heart rate, similar to the results seen in Figure 5a for hybrid optical gating experiments. We observed no developmental abnormalities in samples imaged in our hybrid optical gating microscope and fish appear to undergo normal development with no obvious delay for fish developing at 23° C.

### 7 Assessment of photobleaching during imaging

For retrospective gating, the same method was followed as above. For convenience and consistency, rather than performing an actual retrospective gating analysis on a high-speed recording of fluorescence images, we used prospective optical gating to trigger synchronized acquisition of the images that were subsequently analyzed and compared against results from hybrid optical gating. However, throughout the *z* scan the laser was turned on continuously, exactly as it would be for retrospective gating acquisition.

To quantify the fluorescence signal level in the *z* stack, an estimate of the global background signal level was made and subtracted from all pixels. The values of all voxels *v*(*x, y, t*) within a volume-of-interest around the heart was summed for each timepoint *t*:

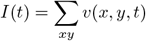

The resultant time-sequence was fitted with a double-exponential *I*(*t*) = *a* + *b*exp(−*ct*) + *d*exp(−*et*). The constant term *a* compensates primarily for faint autofluorescence of red blood cells within the heart: due to the constant turnover of blood cells within the light sheet, this signal is not significantly bleached even at extremely high laser doses. We found that the double-exponential provided a good description of the complex dynamics involved in photobleaching, for the time-sequence from the retrospective gating protocol. A single-exponential was fitted in the case of the hybrid gating protocol, since we found that the time constant of any second exponential term was too long to be reliably determined.

### 8 Cardiac laser-injury

A Zeiss Photo Activated Laser Microdissection microscope system was used to induce a localised laser injury to the ventricular apex of anaesthetised 72 hpf embryos. Embryos were laterally mounted on a glass slide in 20 *μ*l of conditioned water and the laser was fired through a 20× objective. Embryos were deemed injured when ventricular contractility considerably decreased and the ventricular apex reduced in size without rupturing the myocardial wall. Uninjured (control) embryos were treated in the same manner up to the point of laser injury, when they were separated and maintained in the same conditions as injured embryos.

### 9 Neutrophil Tracking in Time-lapse Experiments

Neutrophil tracking was performed using the automated algorithms available in FIJI [46] via the TrackMate plug-in [47]. Neutrophils were detected with an expected diameter of 12 microns and with a “Quality” threshold of 20.0; approximately the same number of neutrophils per frame were detected for all time intervals (Supplementary Figure 2c). Tracks were then recovered with the Simple Linear Assignment Problem algorithms. At three minute time-lapse intervals, a maximum linking distance of 30 microns was used, and a gap-closing maximum distance of 60 microns and 2 frames was used. These settings accommodate a maximum neutrophil velocity of 10 microns/minute. Detections for all time intervals (see below) and tracks at 3 minute time intervals were manually inspected to check there were no obvious errors.

To demonstrate the need for time-lapse intervals as short as 3 minutes, we examined how track features change with increasing time-lapse intervals. First we created two track sets:

1. “Subsampled” virtual experiments were performed by starting with 3 minute time-lapse dataset but retaining only every second stack for a 6 minute interval, every third stack for a 9 minute interval, etc. Tracks were recovered for these subsampled experiments as above, except the linking maximum distance and gap-closing maximum distance were scaled to allow the same maximum neutrophil velocity to be captured (Supplementary Figure 2b shows an example at 30 minute intervals). This track set represents our own best attempt at recovering tracks from data taken at longer time intervals, using off-the-shelf analysis tools.
2. Starting from tracks recovered from the 3 minute time-lapse dataset, “decimated” tracks were created by only keeping track points associated with even-numbered stacks for a 6 minute interval, and so on. Note that these decimated tracks are the closest to a “ground truth” dataset that can be recovered; it is intuitively apparent that the tracks captured at 3 minute will have the fewest accidental track breakages or incorrect linkages (see Figure 4f for an example at 30 minute intervals). This track set therefore represents an upper bound on achievable performance, in which the track analysis was actually performed with additional information (3 minute interval image stacks) available to ensure correct track linking, which would not in reality be available in an experiment using longer time-lapse intervals.

From these two track sets we measured the number of neutrophils per frame as a control (Supplementary Figure 2c), the number of tracks, and the number of tracks that pass through the wound area (Figure 4e). We also measured the average absolute velocity and the meandering index [22] for each individual track (Figure 4f, Supplementary Figure 2c).

### 10 Image acquisition, processing and analysis

Imaging parameters used for all experiments can be found in Supplementary Table 3.

Volume datasets are presented as maximum intensity projections (MIPs) unless otherwise stated. A trivial linear colour mapping, starting at zero, was used in all cases. For the local detail presented in Figures 4 and 6, MIPs were computed over a cropped z range to exclude background clutter. For time series movies (apart from Supplementary Video 9), a small exponential correction was applied as a function of time, to compensate for gradual photobleaching.

Surface rendering was performed using the Imaris software (Bitplane AG). Percentage increase in endothelial chamber volume was calculated by manually cropping the rendered atrium and ventricle and exporting the volume at 48hpf and 72hpf.

### 11 Software and Data handling

The computer code for real-time synchronization, long-term updating and image acquisition is implemented on a unix-like platform (iMac 2015, Apple Inc.) to ensure consistent real-time performance and to benefit from high quality performance diagnostic tools. The real-time computational demands of the analysis allow it to be run on a standard desktop computer, and we have even achieved good results using low-end portable computers (iBook 2008, Apple Inc.). The computer code implements the algorithms described above, and presents a graphical user interface for imaging, microscope control and initial post-experiment visualization and processing (Supplementary Figure 7). In future we intend to migrate our now-mature synchronization system to a cross-platform framework such as Micromanager.

Data is streamed to the computer’s hard disk in real-time. The storage requirements are comparatively modest (approx 100 MB/channel/3D timepoint) since the only fluorescence images acquired and stored are from the desired phase in the heartbeat. This contrasts with alternative post-acquisition approaches [12, 13] that typically require several hundred times this number of images to be recorded for subsequent postprocessing. Maximum intensity projection images of the z-stacks are available for monitoring during acquisition, and on experiment completion data is prepared in a format suitable for importing into visualization software such as Imaris (Bitplane AG).

### 12 Code Availability

The graphical user interface and all codes required to run hybrid optical gating live have been deposited in the University of Glasgow data repository at http://dx.doi.org/10.5525/gla.researchdata.729. This includes codes for prospective optical gating, updating reference frames and the codes needed for hardware interactions. Also available are Python/Jupyter notebooks for recreating the plots in Figures 3–5 and Supplementary Figures 1–2, and also for generating Supplementary Video 2.

### 13 Data Availability

The data used to produce all results in this paper have been deposited in the University of Glasgow data repository at http://dx.doi.org/10.5525/gla.researchdata.729. This repository contains maximum intensity projections of the fluorescence data shown in: Figures 3–6; Supplementary Figures 2 and 4; and Supplementary Videos 4–7 and 9–10. Brightfield reference heartbeats used in Figures 3 and 5, and Supplementary Videos 2 and 8, are also available in this repository. Raw datasets for the experiments in this paper are very large, even with hybrid optical gating, and so most of these are not included in the repository but are available on reasonable request from the corresponding author. However, the full raw data for a small illustrative time range from Figure 6 have been included in this repository. The repository also includes some processed data for the Jupyter Notebooks provided; this has been described in these Notebooks.

## Supplementary Information

**Table 1:** The following table contains the key parameters used in the experiments for Figure 5a. Definitions match those given in [24].

| Parameter | Units | Hybrid Optical Gating | Retrospective Optical Gating<sup>1</sup> |
| --- | --- | --- | --- |
| Wavelength | nm |  | 488 |
| Illumination Time (per individual frame) | ms |  | 2.5 |
| Dark Interval (between frames) | ms | 400<sup>2</sup> | 0<sup>3</sup> |
| Stack Time | s | 82<sup>2</sup> | 225 |
| Dark Interval (between stacks) | s | 200<sup>2</sup> | 75 |
| Number of Images per Plane |  | 1 | 600 |
| Number of Planes per Stack<sup>4</sup> |  |  | 150 |
| Number of Stacks |  |  | 20 |
| Distance between $z$ planes | $\mu\text{m}$ | | 1 |
| Excitation power<sup>5</sup> | mW |  | 1.25 |
| Intensity<sup>5</sup> | $\text{Wcm}^{-2}$ | | 118 |
| Exposure | $\text{Jcm}^{-2}$ | 0.3 | 450 |
| Energy per stack | J | 0.00019 | 0.28 |

<sup>1</sup> Parameters selected to give a light dose similar to that used in [13].
<sup>2</sup> The minimum time between individual slice captures is one heartbeat (approximately 140 beats per minute) but this interval will vary with age and health and photoresponse. This will also affect the total time to collect a whole stack.
<sup>3</sup> The retrospective optical gating protocol uses constant laser illumination for back-to-back image capture.
<sup>4</sup> The size of images and number of images per stack were varied depending on the orientation and size of individual hearts.
<sup>5</sup> We measured our light sheet to be approximately $2.4 \mu\text{m}$ thick (FWHM) and $390 \mu\text{m}$ tall (FWHM) at its centre. We measured that approximately 11% of the power configured in the laser software arrived at the sample, after optical losses.

**Table 2:** The following table contains the key parameters used in the experiments for Figure 5b and Supplementary Video 9. Definitions match those given in [24].

| Parameter | Units | Hybrid Optical Gating | Retrospective Optical Gating<sup>1</sup> |
| --- | --- | --- | --- |
| Wavelength | nm |  | 561 |
| Illumination Time (per individual frame) | ms |  | 15 |
| Dark Interval (between frames) | ms | 400<sup>2</sup> | 0<sup>3</sup> |
| Stack Time | s | 60<sup>2</sup> | 240 |
| Dark Interval (between stacks) | s | 240<sup>2</sup> | 60 |
| Number of Images per Plane |  | 1 | 106 |
| Number of Planes per Stack<sup>4</sup> |  | 140 | 150 |
| Number of Stacks |  | 262 | 29 |
| Distance between $z$ planes | $\mu\text{m}$ | | 1 |
| Excitation power<sup>5</sup> | mW |  | 1.2 |
| Intensity<sup>5</sup> | $\text{Wcm}^{-2}$ | | 113 |
| Exposure | $\text{Jcm}^{-2}$ | 4.3 | 460 |
| Energy per stack | J | 0.0025 | 0.29 |

<sup>1</sup> Parameters selected to give a light dose similar to that used in [13].
<sup>2</sup> The minimum time between individual slice captures is one heartbeat (approximately 140 beats per minute) but this interval will vary with age and health and photoresponse. This will also affect the total time to collect a whole stack.
<sup>3</sup> The retrospective optical gating protocol uses constant laser illumination for back-to-back image capture.
<sup>4</sup> The size of images and number of images per stack were varied depending on the orientation and size of individual hearts. For this reason, the number of planes is slightly different between the two fish here.
<sup>5</sup> We measured our light sheet to be approximately $2.4 \mu\text{m}$ thick (FWHM) and $390 \mu\text{m}$ tall (FWHM) at its centre. We measured that approximately 11% of the power configured in the laser software arrived at the sample, after optical losses.

**Table 3:** The following table contains the key parameters used in the experiments for all figures. Definitions match those given in [24].

| Parameter | Units | Figure |  |  |  |
| --- | --- | --- | --- | --- | --- |
|  |  | 3a (Supp. Video 4) | 4a (Supp. Video 5) | 4b (Supp. Figure 4, Video 6) | 6 (Supp. Figure 5, Video 7) |
| Fluorophore: Wavelength | nm | mCherry:561 | GFP:488; mCherry:561 | GFP:488; mCherry:561 | GFP:488; mKate:561 |
| Illumination Time (per individual frame) | ms | 30 | 7; 15 | 7; 30 | 10; 15 |
| Dark Interval/ms (between frames)<sup>1</sup> | ms | 470 | 360 | 470 | 400 |
| Stack Time/s<sup>1</sup> | s | 110 | 50 | 32 | 60 |
| Dark Interval/s (between stacks)<sup>1</sup> | s | 190 | 130 | 148 | 240 |
| Number of Images per Plane |  | 1 | 1 | 1 | 1 |
| Number of Planes per Stack |  | 220 | 131 | 64 | 140 |
| Number of Stacks |  | 288 | 200 | 460 | 262 |
| Distance between $z$ planes | $\mu\text{m}$ | 1 | 0.75 | 1.5 | 1 |
| Excitation power<sup>2</sup> | mW | 11 | 11 | 11 | 11 |
| Intensity<sup>2</sup> | $\text{Wcm}^{-2}$ | 1010 | 1010 | 1010 | 1010 |
| Exposure | $\text{Jcm}^{-2}$ | 79 | 19; 40 | 19; 79 | 26; 40 |
| Energy per stack | J | 0.073 | 0.010; 0.022 | 0.010; 0.043 | 0.015; 0.023 |

<sup>1</sup> Numbers are approximate. The minimum time between individual slice captures is one heartbeat (approximately 140 beats per minute; 155 beats per minute for figure 4a) but this interval will vary with age and health. This will also affect the total time to collect a whole stack.
<sup>2</sup> We measured our light sheet to be approximately $2.4 \mu\text{m}$ thick (FWHM) and $390 \mu\text{m}$ tall (FWHM) at its centre. We measured that approximately 11% of the power configured in the laser software arrived at the sample, after optical losses.

**Supplementary Figure 1:**
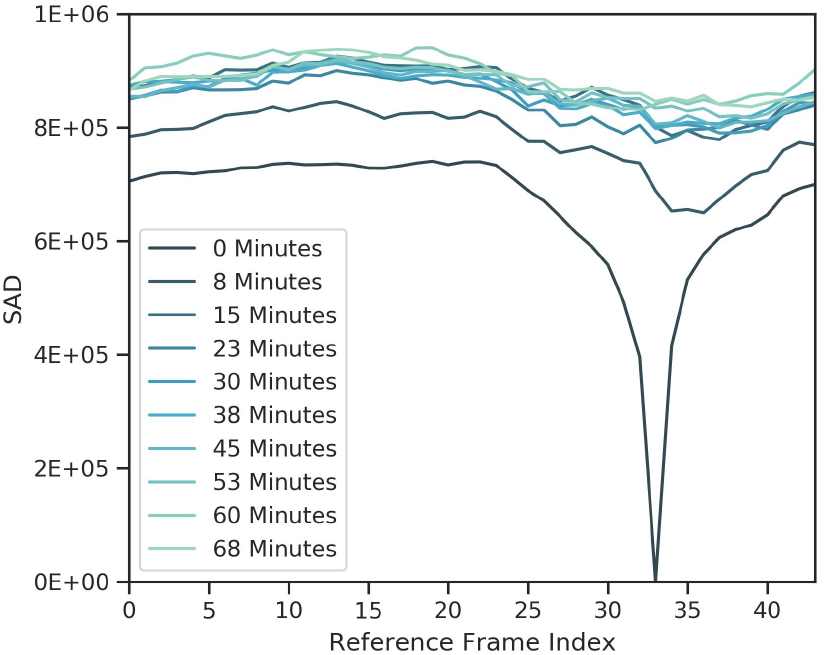
Over time, prospective optical gating loses certainty of phase recovery, preventing long-term imaging. As time progresses (colour gradient) the shape of the sum-of-absolute-differences (SAD) comparison with the reference heartbeat changes, losing the clear, sharp peak needed for accurate phase recovery. This can happen after as little as 20 minutes. By 60 minutes or more, prospective optical gating is no longer able to accurately phase-match the incoming brightfield images (see supplementary video 2) and hybrid optical gating methods are required to update the reference heartbeat (see figure 2).

**Supplementary Figure 2:**
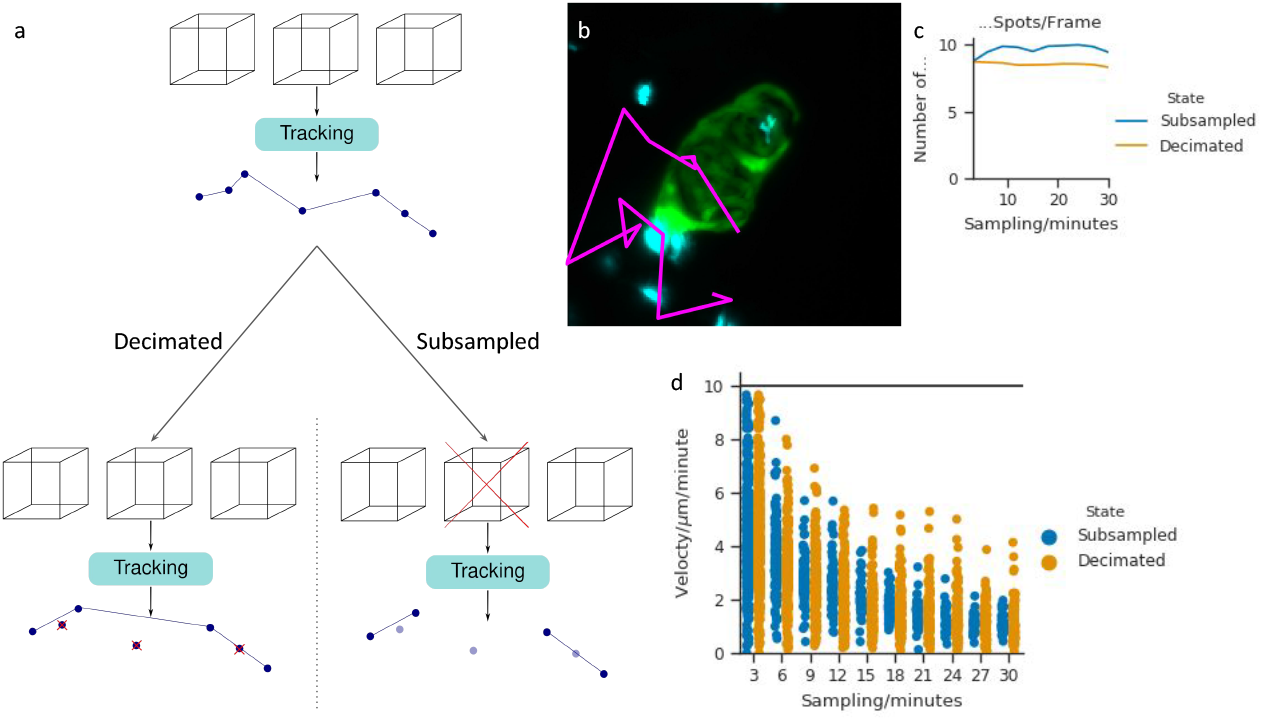
Hybrid optical gating time-lapse experiments allow accurate cell tracking compared to time-lapse intervals possible with other 3D approaches. **a,** In order to produce ‘ground truth’ tracks with different sampling intervals we recovered tracks at a 3 minute sampling rate and *decimated* these recovered tracks to produce tracks with true connections. We compared this to tracks recovered from *subsampled* time-lapse data, **b,** Cell tracking in time-lapse data with large time intervals, here 30 minutes, can lead to very short tracks (not shown) due to the large movement of cells between images, or erroneous and erratic paths (example in pink). Fluorescent labelling shows cardiomyocytes (green - transgene *myl7:GFP*) and neutrophils (cyan - transgene *mpeg1:mCherry*). **c,** Number of detections per frame when comparing subsampled image data (blue) against ‘ground truth’ decimated tracks (orange). This acts as a control that the decimated and subsampled data are using similar detected nuclei, **d,** Track errors at higher sampling intervals cause quantification errors when trying to assess neutrophil behaviour. Here we show the average absolute velocity for tracks recovered from subsampled image data (blue) and decimated tracks (orange); the solid line shows the theoretical maximum velocity for this analysis. In both cases very fast neutrophils are lost and the average velocity across the tracks drops; however, in the subsampled case, erroneous track recovery also loses tracks with very low velocity as these are incorrectly merged with other tracks.

**Supplementary Figure 3:**
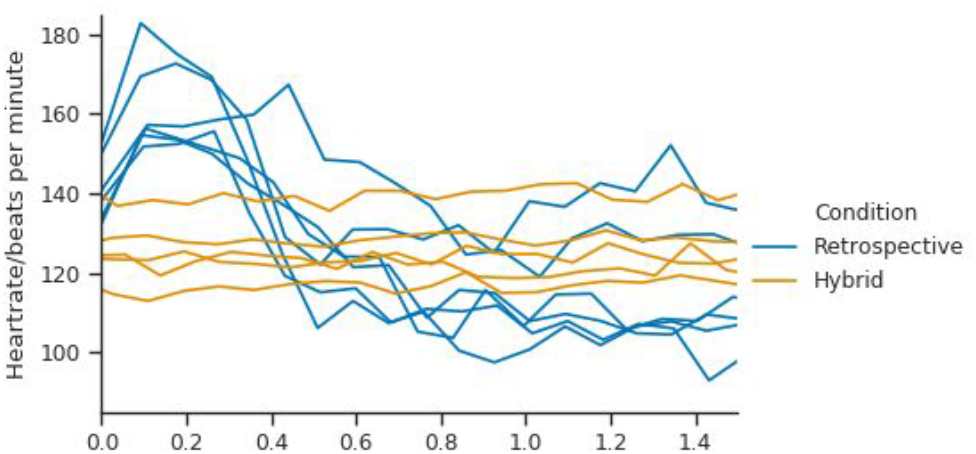
Hybrid optical gating protocol causes less cardiac photo-injury and photo-bleaching than retrospective gating protocols. A replotting of Figure 5a showing individual fish.

**Supplementary Figure 4:**
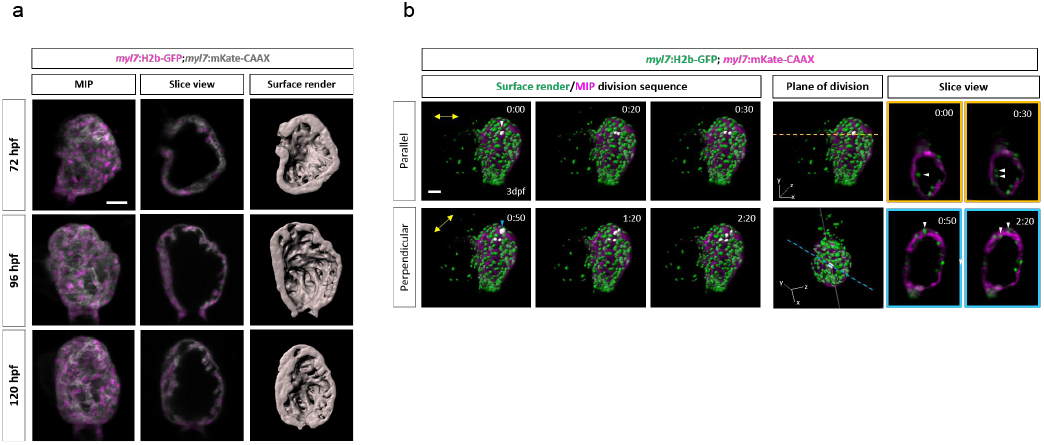
Hybrid optical-gating facilitates examination of trabecular development at the tissue and cellular level. **a,** A single *myl7:h2b-GFP;myl7:mKate-CAAX* ventricle imaged in three discrete 6 hour time-lapses beginning at 3dpf, 4 dpf and 5dpf. Representative images from each time-lapse of this heart are shown as a MIP (left), midpoint slice (middle) and luminal surface render showing the developing trabecular network (right). Cardiomyocyte nuclei are shown in magenta and cardiomyocyte membranes in gray. **b,** Representative timepoints from a timelapse of whole 3dpf heart showing surface-rendered cardiomyocyte nuclei (green - transgene *myl7:h2b-GFP*) superimposed onto a MIP of cardiomyocyte membranes (magenta - transgene *myl7:mKate-CAAX*). The white arrowhead indicates a cardiomyocyte nucleus prior to its division parallel to the myocardial wall (top row). Blue arrowhead indicates a cardiomyocyte nucleus prior to a division, perpendicular to the myocardial wall (bottom row). The slice view shows each dividing nuclei pre and post-division in the plane of division where dividing nuclei are indicated by a white arrow head. By studying the pre and post-division images in the slice view, it becomes clear that one nucleus has divided parallel to the myocardial wall (top) and the other has divided approximately to it (bottom). The plane of division is shown as a yellow or blue dashed line for the parallel and perpendicular division respectively. Scale bars: 30 *μ*m.

**Supplementary Figure 5:**
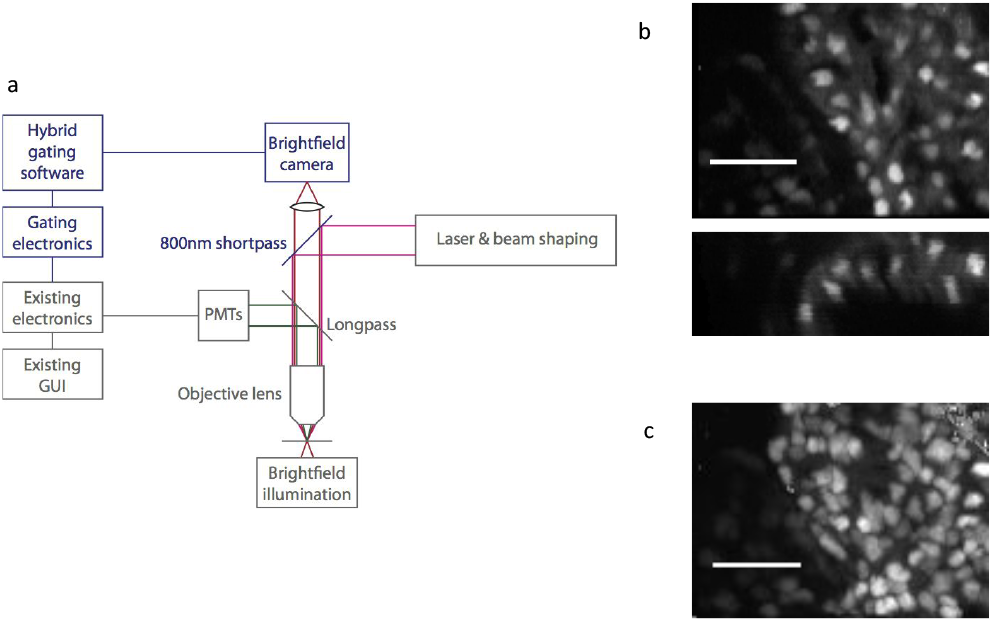
Synchronized two-photon microscopy. **a,** Schematic diagram of Scientifica commercial two-photon microscope (grey) and the minor modifications (black) required to integrate our prospective optical gating system. **b, c,** xy and xz projections (**b**) and maximum intensity projection (c) of synchronized two-photon z stack, showing cardiomyocyte nuclei (transgene *myl7:h2b-gfp*) in region of interest in the zebrafish ventricle (5 dpf). Scale bars: 30 *μ*m.

**Supplementary Figure 6:**
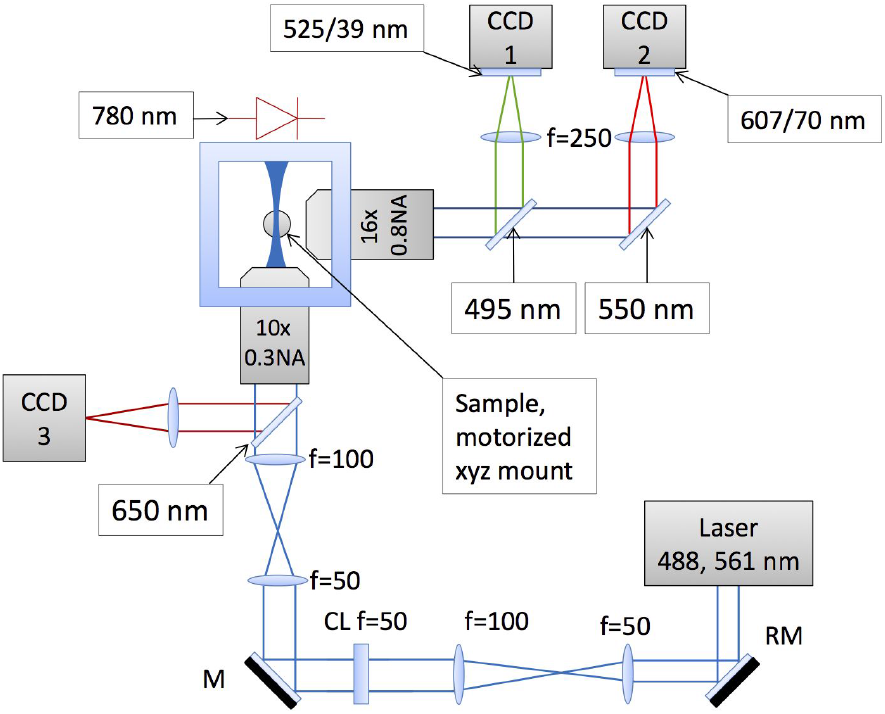
Optical setup for microscope used to demonstrate hybrid optical gating. A light sheet is launched into the sample chamber via a water-dipping objective lens. The same objective lens is simultaneously used for brightfield near-infrared imaging (CCD3). A separate water-dipping objective lens is used for imaging fluorescence in multiple colour channels (CCD1, 2). The hybrid optical gating algorithms analyze the brightfield images from CCD3, and generate trigger signals that cause synchronized acquisition of individual fluorescence frames by CCD1 and CCD2. Lasers are pulsed such that they only illuminate the sample during the exposure window of the relevant CCD camera.

**Supplementary Figure 7:**
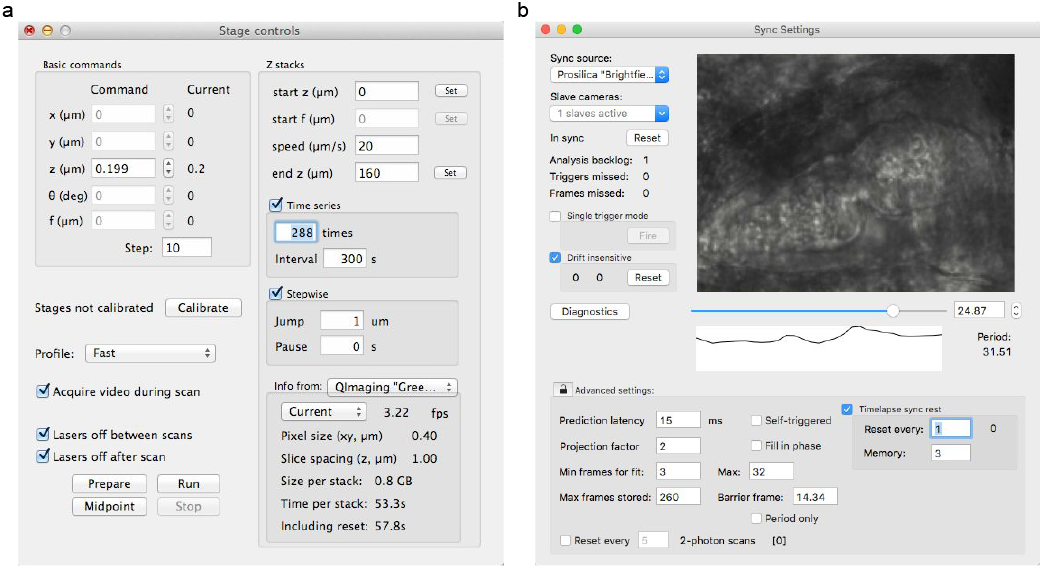
User interface for hybrid optical gating. Example screenshots from our complete graphical user interface for timelapse optically-gated imaging experiments. **a,** Timelapse z stacks are programmed to repeat at regular intervals, with uniformly-spaced z slices acquired at a fixed phase in the cardiac cycle. **b,** Hybrid optical gating is configured by the user at the start of the experiment, and it then runs without further user intervention. Key features include: selection of ‘slave’ cameras for triggered acquisition; selection of target phase in the cardiac cycle; regular updates of the image sequence representing our reference heartbeat (‘timelapse sync reset’) in order to adapt to the changing appearance of the heart over developmental timescales.

## Supplementary Note 1: invariant brightfield source images

Any image-based heart synchronization strategy performs best when the appearance of the heart is invariant over time (aside from periodic changes over the heartbeat). In synchronized 3D imaging this gives rise to a conflict between the need to scan the fluorescence image plane through the sample (or vice-versa) for 3D imaging, and the desire to image a fixed plane in order to obtain a consistent phase reference for synchronization. Various strategies for dealing with this issue can be found in the literature:

- Attempting to maintain phase-lock despite gradual changes in appearance from one slice to the next [12] (but this approach suffers from accumulated systematic error in z [13, 48])
- Refocusing the brightfield images using a motorized mobile tube lens [11] (but this is cumbersome and in practice limits acquisition to a z range of 120 *μ*m).
- Use of a rotational imaging strategy, with synchronization being based on a line of pixels along the axis of rotation [48].

In our present work we instead acquire our phase-reference images via brightfield imaging through the light sheet launch objective. Consequently, during a z scan the brightfield images remain focused on the same plane in the sample, but the image translates within the focal plane of the brightfield camera. Our heart synchronisation algorithms are already tolerant of such in-plane motion (to accommodate possible sample drift), and thus we are able to maintain a high-quality phase-lock throughout a whole z scan.

When imaging on a commercial microscope (for example the two-photon imaging in Supplementary Figure 5) it is not always possible to use side view imaging or focus correction. Under those circumstances it is necessary to fall back on strategies where the reference frames are intermittently updated as the appearance of the heart in the brightfield channel changes during a z scan (a strategy originally presented in [12] for retrospective synchronization). In our case we apply exactly the same reference-frame-refresh strategy that we use for longterm time-lapse imaging, but here we force a refresh after every third two-photon image acquisition over the course of a z scan.

## Supplementary Note 2: hardware interfacing

### Timing hardware

Typical desktop computers do not have a standardised interface for simple digital input/output, suitable for communicating with hardwares such as cameras. Furthermore, it is not trivial to write code with guaranteed predictability to enable millisecond-resolution accuracy in hardware interactions. For these reasons, our platform is design to interface over an RS-232 serial link to a custom-built microcontroller system that generates the actual electrical trigger signals at the times determined by our synchronization software. In practice, this timing hardware fulfils other roles as well, pulsing lasers in time with camera exposures, generating custom digital waveforms for particle image velocimetry, etc. A simple text-based communications protocol is defined (see document “communications protocol.rtf” in supplementary source code) for communication between the main computer software and the timing hardware.

Source code and schematics for our custom timing hardware have been made available, but any hardware conforming to the specified communications protocol could be used. Other hardware with compatible capabilities could be used by adapting our source code (new subclass of TimingBoxBase, modelling code on existing subclasses TimingBoxXMOS and TimingBoxFPGA) to support a different communications protocol.

### Camera support

Currently our user interface supports the following camera models:

- Prosilica (recommended as brightfield source for heart synchronization) via PvAPI driver for GigE cameras
- Ximea (XiD, XiQ) via xiAPI
- QImaging via deprecated QCam API

Other camera hardware and drivers can be supported by subclassing CameraBase/FrameBase classes, modelling code on the existing camera support classes.

### Translation stage support

Currently our user interface supports the following translation stage models:

- Physik Instrumente via the Mercury native command interface
- Newport via SMC100 command set

Other stage hardware can be supported by subclassing the StageCommandInterface class, modelling code on existing stage support classes.

## Supplementary Video Captions

The following Supplementary Videos are available alongside this preprint.

Supplementary Video 1: **Optical gating concept.** The constant motion of the beating heart presents a challenge for imaging (grey representation of part of the heart ventricle). Light sheet fluorescence microscopy provides optical sectioning to image a single plane of the heart at a time (depicted in green), without any superfluous illumination in other spatial locations. However, retrospective optical gating techniques require many raw fluorescence video images to be acquired at *every* z plane in the heart; a consistent z stack at a fixed cardiac phase is reconstructed following subsequent computer analysis. In contrast, prospective optical gating ensures that the heart is only exposed to potentially-harmful laser light at the required *time* in the cardiac cycle.

Supplementary Video 2: **Prospective optical gating fails to maintain phase-lock over extend periods.** Due to changes in brightfield features, prospective optical gating loses phase-lock with time (left) whilst hybrid optical gating is able to maintain a lock (right) over many hours. The colours indicate the difference in phase between the frame shown and the mean human-scored target frame (c.f. Figure 3).

Supplementary Video 3: **Diagrammatic representation of hybrid optical gating for time-lapse 3D imaging of the beating zebrafish.** For each time-lapse point, a new reference heartbeat is determined in the brightfield images. This new reference heartbeat could be used to create a 3D stack using prospective optical gating; however, the new reference heartbeat is not aligned with previous reference sets (blue). By incorporating retrospective optical gating on these brightfield reference heart beats, including motion correction and a weighted regression with historical reference sets, the previous target can be identified in each new reference set. This allows phase-locking between stacks throughout a time-lapse experiment (right).

Supplementary Video 4: **Hybrid optical gating allows phase-locked, longterm, time-lapse imaging and quantification of developmental cardiac morphogenesis.** Phase-locked, live time lapse imaging of heart development (48 to 72 hours post-fertilization (hpf), at 300s intervals) showing endothelium lining blood vessels and heart chambers (transgene *flk1:mCherry*). Movie shows maximum intensity projections of z-stacks. Scale bar: 30*μ*m.

Supplementary Video 5: **Hybrid optical gating allows phase-locked, longterm, time-lapse imaging and quantification of developmental cardiac morphogenesis.** Phase-locked, live time lapse imaging of heart development (48 to 72 hours post-fertilization (hpf), at 300s intervals) showing endothelium lining blood vessels and heart chambers (transgene *flk1:mCherry*). Movie shows 3D rendering of heart chambers. Scale bar: 30*μ*m.

Supplementary Video 6: **Hybrid optical gating allows phase-locked time-lapse imaging at the rate required for investigating cell activity during injury response.** Synchronized time-lapse 3D imaging of immune cells migrating to the site of laser-injury in a zebrafish heart. Maximum intensity projection of z-stacks, showing macrophages (magenta - transgene *mpeg:mCherry*) at the injury site on cardiomyocytes (green - transgene *myl7:GFP*). Injury occurred at 74 hpf, with imaging commencing at approx 83.5 hpf; note that the effect of laser injury on heart rhythm posed challenges for the synchronization, resulting in some planes and/or timepoints with poor synchronization. Scale bar: 30*μ*m.

Supplementary Video 7: **Hybrid optical gating allows phase-locked time-lapse imaging at the rate required for investigating cell activity during injury response.** Synchronized time-lapse 3D imaging of neutrophil immune cells migrating to the site of laser-injury in a zebrafish heart. Maximum intensity projection of z-stacks, showing neutrophils (cyan - transgene *mpx:mCherry*) at the injury site on the apex of the ventricle (green - transgene *myl7:GFP*). Injury was induced at 72 hpf, with imaging commencing at approx 74 hpf. Scale bar: 30*μ*m.

Supplementary Video 8: **Retrospective gating affects heart function.** We have observed changes in heart function when using retrospective gating protocols to acquire timelapse datasets. Video shows visible deterioration of ejection fraction, heart rate and rhythm.

Supplementary Video 9: **Hybrid optical gating substantially reduces bleaching rate in sensitive samples.** When fluorophores are particularly light-sensitive, or labelling is highly specific and therefore sparse (e.g. *myl7:mKate-CAAX* transgenic line), we observe that specimens are easily bleached, even using light sheet fluorescence microscopy. When exposed to light levels required for retrospective gating (using typical parameters reported in the literature), the signal from a specimen labelled with membrane-specific mKate fluorophore is halved after only 3 timepoints, whereas with hybrid optical gating over 150 timepoints are recorded before the signal level drops by the same amount.

Supplementary Video 10: **3D time-lapse imaging of cardiomyocyte migration and cell division during cardiac trabeculation.** 24 hour timelapse (shown here as a Maximum Intensity Projection) of a 72-96hpf embryo, showing cardiomyocyte nuclei (green - transgene *myl7:h2b-gfp*) and cardiomyocyte membranes (magenta - transgene *myl7:mKate-CAAX*). Nuclei of cardiomyocytes which are about to divide are marked by white arrowheads, or open-arrowheads (less-obvious divisions near the posterior wall of the heart).

## References

[1] Paul J Scherz et al. “High-speed imaging of developing heart valves reveals interplay of morphogenesis and function”, Development (Cambridge, England) 135 (2008), pp. 1179–87. DOI: 10.1242/dev.010694.

[2] Philipp J Keller et al. “Fast, high-contrast imaging of animal development with scanned light sheet-based structured-illumination microscopy”, Nature methods 7 (2010), pp. 637–642. DOI: 10.1038/nmeth.1476.

[3] Nicolas Olivier et al. “Cell lineage reconstruction of early zebrafish embryos using label-free nonlinear microscopy”, Science (New York, N.Y.) 329 (2010), pp. 967–71. DOI: 10.1126/science.1189428.

[4] Jim Swoger et al. “4D retrospective lineage tracing using SPIM for zebrafish organogenesis studies”, Journal of biophotonics 4 (2011), pp. 122–34. DOI: 10.1002/jbio. 201000087.

[5] David W Staudt et al. “High-resolution imaging of cardiomyocyte behavior reveals two distinct steps in ventricular trabeculation”, Development (Cambridge, England) 141 (2014), pp. 585–93. DOI: 10.1242/dev.098632.

[6] Veronica Uribe et al. “In vivo analysis of cardiomyocyte proliferation during trabeculation”, Development 145 (2018), dev164194.

[7] K. Greger, J. Swoger, and E. H. K. Stelzer. “Basic building units and properties of a fluorescence single plane illumination microscope”, Review of Scientific Instruments 78 (2007), p. 023705. DOI: 10.1063/1.2428277.

[8] Rory M Power and Jan Huisken. “A guide to light-sheet fluorescence microscopy for multiscale imaging”, Nature Methods 14 (2017), pp. 360–373. DOI: 10.1038/nmeth.4224.

[9] Michael Liebling et al. “Rapid three-dimensional imaging and analysis of the beating embryonic heart reveals functional changes during development”, Developmental dynamics 235 (2006), pp. 2940–8. DOI: 10.1002/dvdy.20926.

[10] Jonathan M. Taylor et al. “Real-time optical gating for three-dimensional beating heart imaging”, Journal of Biomedical Optics 16 (2011), p. 116021. DOI: 10.1117/1.3652892.

[11] Jonathan M Taylor, John M Girkin, and Gordon D Love. “High-resolution 3D optical microscopy inside the beating zebrafish heart using prospective optical gating”, Biomedical optics express 3 (2012), pp. 3043–53. DOI: 10.1364/BOE.3.003043.

[12] Michael Liebling et al. “Four-dimensional cardiac imaging in living embryos via postacquisition synchronization of nongated slice sequences”, Journal of Biomedical Optics 10 (2005), p. 054001. DOI: 10.1117/1.2061567.

[13] Michaela Mickoleit et al. “High-resolution reconstruction of the beating zebrafish heart”, Nature methods 11 (2014), pp. 1–6. DOI: 10.1038/nmeth.3037.

[14] Michael Weber et al. “Cell-accurate optical mapping across the entire developing heart”, eLife 6 (2017), pp. 1–14. DOI: 10.7554/eLife.28307.

[15] Jungho Ohn, Huai-Jen Tsai, and Michael Liebling. “Joint dynamic imaging of morphogenesis and function in the developing heart”, Organogenesis 5 (2009), pp. 248–55.

[16] Jenny Pestel et al. “Real-time 3D visualization of cellular rearrangements during cardiac valve formation”, Development 143 (2016), pp. 2217–2227. DOI: 10.1242/dev.133272.

[17] Martin A Denvir, Carl S Tucker, and John J Mullins. “Systolic and diastolic ventricular function in zebrafish embryos: influence of norepenephrine, MS-222 and temperature”, BMC biotechnology 8 (2008), p. 21.

[18] Peter J. Rombough. “Ontogenetic changes in the toxicity and efficacy of the anaesthetic MS222 (tricaine methanesulfonate) in zebrafish (Danio rerio) larvae”, Comparative Biochemistry and Physiology Part A: Molecular & Integrative Physiology 148 (2007), pp. 463–469. DOI: https://doi.org/10.1016/j.cbpa.2007.06.415.

[19] DY Stainier, Robert K Lee, and Mark C Fishman. “Cardiovascular development in the zebrafish. I. Myocardial fate map and heart tube formation”, Development 119 (1993), pp. 31–40.

[20] Stefan Rohr, Cecile Otten, and Salim Abdelilah-Seyfried. “Asymmetric involution of the myocardial field drives heart tube formation in zebrafish”, Circulation Research 102 (2008), e12–e19.

[21] Dimitris Beis et al. “Genetic and cellular analyses of zebrafish atrioventricular cushion and valve development”, Development 132 (2005), pp. 4193–4204.

[22] Katherine M Henry et al. “PhagoSight: an open-source MATLAB@ package for the analysis of fluorescent neutrophil and macrophage migration in a zebrafish model”, PloS one 8 (2013), e72636.

[23] Gianfranco Matrone et al. “Laser-targeted ablation of the zebrafish embryonic ventricle: A novel model of cardiac injury and repair”, International Journal of Cardiology 168 (2013), pp. 3913–3919.

[24] P Philippe Laissue et al. “Assessing phototoxicity in live fluorescence imaging”, Nature Methods 14 (2017), pp. 657–661. DOI: 10.1038/nmeth.4344.

[25] Jaroslav Icha et al. “Phototoxicity in live fluorescence microscopy, and how to avoid it”, Bioessays 39 (2017), p. 1700003.

[26] Omar E. Olarte et al. “Light-sheet microscopy: a tutorial”, Adv. Opt. Photon. 10 (2018), pp. 111–179. DOI: 10.1364/A0P.10.000111.

[27] Loling Song et al. “Photobleaching kinetics of fluorescein in quantitative fluorescence microscopy”, Biophysical journal 68 (1995), pp. 2588–2600.

[28] Paula J Cranfill et al. “Quantitative assessment of fluorescent proteins”, Nature methods 13 (2016), p. 557.

[29] Alexey M Bogdanov et al. “Cell culture medium affects GFP photostability: a solution”, nature methods 6 (2009), p. 859.

[30] Jiandong Liu et al. “A dual role for ErbB2 signaling in cardiac trabeculation”, Development 137 (2010), pp. 3867–3875.

[31] Jonathan M Taylor. “Optically gated beating-heart imaging”, Frontiers in Physiology 5 (2014), p. 481. DOI: 10.3389/fphys.2014.00481.

[32] Arian S Forouhar et al. “The embryonic vertebrate heart tube is a dynamic suction pump”, Science 312 (2006), pp. 751–3. DOI: 10.1126/science.1123775.

[33] V. Zickus and J. M. Taylor. “3D + time blood flow mapping using SPIM-microPIV in the developing zebrafish heart”, Biomedical Optics Express 9 (2018), pp. 2418–2435.

[34] Hang Yu et al. “Combining Near-infrared Excitation with Swept Confocally-aligned Planar Excitation (SCAPE) Microscopy for Fast, Volumetric Imaging in Mouse Brain”, Biophotonics Congress: Biomedical Optics Congress 2018. Optical Society of America, 2018, BF3C.3. DOI: 10.1364/BRAIN.2018.BF3C.3.

[35] Omar E. Olarte et al. “Decoupled illumination detection in light sheet microscopy for fast volumetric imaging”, Optica 2 (2015), p. 702. DOI: 10.1364/OPTICA.2.000702.

[36] Thai V. Truong et al. “Selective volume illumination microscopy offers synchronous volumetric imaging with high contrast”, bioRxiv (2018). DOI: 10.1101/403303. eprint: https://www.biorxiv.org/content/early/2018/08/29/403303.full.pdf.

[37] Vikas Gupta and Kenneth D Poss. “Clonally dominant cardiomyocytes direct heart morphogenesis”, Nature 484 (2012), p. 479.

## References

[38] M Westerfield. The zebrafish book: a guide for the laboratory use of zebrafish (Danio rerio). Univ. of Oregon Press, Eugene, 2000.

[39] C Geoffrey Burns et al. “High-throughput assay for small molecules that modulate zebrafish embryonic heart rate”, Nature chemical biology 1 (2005), p. 263.

[40] Kira Proulx, Annie Lu, and Saulius Sumanas. “Cranial vasculature in zebrafish forms by angioblast cluster-derived angiogenesis”, Developmental biology 348 (2010), pp. 3446.

[41] Sa Kan Yoo et al. “Differential regulation of protrusion and polarity by PI (3) K during neutrophil motility in live zebrafish”, Developmental cell 18 (2010), pp. 226–236.

[42] Felix Ellett et al. “mpeg1 promoter transgenes direct macrophage-lineage expression in zebrafish”, Blood 117 (2011), e49–e56.

[43] Yi-Fan Lin, Ian Swinburne, and Deborah Yelon. “Multiple influences of blood flow on cardiomyocyte hypertrophy in the embryonic zebrafish heart”, Developmental biology 362 (2012), pp. 242–253.

[44] J Huisken and D Y R Stainier. “Even fluorescence excitation by multidirectional selective plane illumination microscopy (mSPIM)”, Opt. Lett. 32 (2007), pp. 2608–2610.

[45] Carl J. Nelson et al. “Imaging the developing heart: synchronized time-lapse microscopy during developmental changes”, Proc. SPIE 10499 (2018), 104991F. DOI: 10.1117/12.2290191.

[46] Johannes Schindelin et al. “Fiji: an open-source platform for biological-image analysis”, Nature methods 9 (2012), p. 676.

[47] Jean-Yves Tinevez et al. “TrackMate: An open and extensible platform for single-particle tracking”, Methods 115 (2017), pp. 80–90.

## References

[48] Irina V Larina et al. “Sequential Turning Acquisition and Reconstruction (STAR) method for four-dimensional imaging of cyclically moving structures”, Biomedical optics express 3 (2012), pp. 650–60. DOI: 10.1364/B0E.3.000650.

